# Repeated listening induces exposure-specific cortical tracking of intelligible continuous speech

**DOI:** 10.64898/2026.08.28.747801

**Authors:** Alexandra K. Emmendorfer, Lars Riecke, Hendrik Kröger, Elizaveta Zavialova, Lars Hausfeld

**Affiliations:** Department of Cognitive Neuroscience, Faculty of Psychology and Neuroscience, Maastricht University; Section Teaching and Innovation of Learning, Faculty of Psychology and Neuroscience, Maastricht University; Maastricht Brain Imaging Centre, Maastricht University; Department of Psychiatry, Psychosomatic Medicine and Psychotherapy, University Hospital, Goethe University, Frankfurt am Main, Germany; School of Computation, Information and Technology, Technical University of Munich

**Keywords:** Cortical speech tracking, prediction, EEG, encoding models, mTRF

## Abstract

Neural encoding of acoustic and linguistic features of continuous speech is sensitive to cognitive factors, such as attention and comprehension. We investigated whether neural tracking is also sensitive to the predictability of speech. Participants were repeatedly exposed to intelligible or unintelligible versions of the same audiobook segment while EEG was recorded. First, we fit encoding models to predict EEG responses from acoustic, sublexical, and lexical features of the presented speech. Model comparisons revealed no reliable improvement in model fit when lexical features were included; subsequent analyses were performed on models including only acoustic and sublexical predictors. Second, we compared prediction accuracy for models trained and tested on the same exposures with models trained and tested across different exposures. While we observed no overall change in prediction performance across exposures, we found that models were exposure-specific: prediction performance was highest within the same exposure and decreased with increasing temporal distance between the training and test exposure. This effect was observed for intelligible but not for unintelligible speech, suggesting that the effect depends on properties unique to intelligible speech, such as the ability to form increasingly specific predictions about upcoming linguistic input, rather than general, non-linguistic factors related to repeated exposure. This distance effect was associated with increased model weights from −90 ms to 130 ms, indicating an enhancement of familiar input during an early cortical processing stage. In summary, these findings indicate that cortical tracking of sublexical speech features is modulated by repeated exposure to intelligible speech, consistent with a role for linguistic predictability.

## 1 Introduction

The natural speech signals we encounter in everyday scenarios are highly complex and noisy, yet we are able to process them in a highly efficient manner. Dominant theories of auditory and language processing propose predictive processes as one mechanism contributing to this efficiency (Ferreira & Chantavarin, 2018; Norris et al., 2016; Pickering & Gambi, 2018). According to the predictive coding framework, the brain represents the deviations between expected and actual input, thereby reducing the processing effort for anticipated events (Aitchison & Lengyel, 2017; Friston, 2005; Rao & Ballard, 1999; Spratling, 2017). Evidence for predictive speech processing has been demonstrated across levels of linguistic analysis and modalities, where more predictable or regular signals are associated with more efficient neural processing, as indexed by reduced amplitudes or shorter latencies. For instance, at the sublexical level our brain is sensitive to regularities in combinations of speech sounds (Bonte et al., 2005; Emmendorfer et al., 2020, 2025; Noordenbos et al., 2013; Vidal et al., 2019) and speech rhythm (Bohn et al., 2013; Emmendorfer et al., 2023; Kotz et al., 2018). At the conceptual level, listeners can make use of prior knowledge, discourse context, and visual information to form predictions about the upcoming utterance (Barthel et al., 2024; Ter Bekke et al., 2025; Trujillo et al., 2025).

Much of this evidence comes from experiments that are tightly controlled and using stimuli that bear little resemblance to the perceptual input we encounter in our day-to-day life. In studies using more naturalistic and continuous speech stimuli, it has been shown that the neural activity in the human cortex reliably follows (or tracks) various acoustic and linguistic stimulus features (Brodbeck & Simon, 2020), such as the stimulus envelope (Haufeld et al., 2024; Issa et al., 2024; Kubanek et al., 2013), phonetic (Brodbeck et al., 2018; Di Liberto et al., 2018; Gwilliams et al., 2022; Lesenfants et al., 2019), or semantic information (Broderick et al., 2018; Carta et al., 2025; Gillis et al., 2021; Koskinen et al., 2020; Weissbart et al., 2020). This cortical speech tracking is sensitive to perceptual (Chen et al., 2023; Etard & Reichenbach, 2019; Vanthornhout et al., 2018), linguistic (Brodbeck et al., 2024; Di Liberto et al., 2021), and cognitive factors such as attention (Carta et al., 2025; Hausfeld et al., 2018; O’Sullivan et al., 2015). By modelling the neural response to linguistic features, such as phoneme or semantic surprisal, previous research has revealed that the brain encodes these features indicating EEG signatures classically associated with predictive speech processing (e.g., N400: Broderick et al., 2018; Carta et al., 2025; Heilbron et al., 2022; Weissbart et al., 2020). However, it remains unclear how increasing predictability of an incoming speech signal as a whole influences the neural tracking of its features, and whether this effect differs across features.

Recently, Schubert, Schmidt and colleagues (2023) demonstrated a relationship between individual prediction tendencies (quantified based on pre-stimulus MEG activity in an entropy modulation paradigm) and cortical speech tracking. The left temporal lobe was found to encode the speech envelope with higher accuracy when the speech contained words with high surprisal compared to words with low surprisal, and this effect increased with increasing individual prediction tendencies. A similar pattern has been observed for music envelope tracking, where stronger tracking was observed for atonal music (i.e., music with low pitch predictability) compared to tonal music (high pitch predictability; Keitel et al., 2025). These results suggest that increased predictability of a signal may be associated with reduced neural tracking, in line with theories of predictive processing (Aitchison & Lengyel, 2017; Friston, 2005; Rao & Ballard, 1999; Spratling, 2017).

In the current experiment, we increased the predictability of a speech signal through repeated exposure by presenting the same 30-s audiobook segment five consecutive times, either as clean speech or single-channel noise-vocoded speech. With each exposure, participants became more familiar with the signal, thereby increasing its predictability. Forward encoding models were trained using predictors for three stimulus feature sets of increasing complexity: acoustic, sublexical and lexical. We first investigated whether neural encoding is modulated by predictability, by comparing model performance across exposures. If increased predictability results in reduced encoding accuracy (Keitel et al., 2025; Schubert, Schmidt et al., 2023), we expect model performance to decrease parametrically with increasing exposures. In additional exploratory analyses, we investigate whether neural encoding of the speech signal is exposure-specific, by testing and comparing the performance of models that were trained on different exposures. Finally, we investigate the time-course of exposure-induced changes in neural encoding, by comparing the temporal response functions (TRFs) for the different exposures.

## 2 Methods

### 2.1 Participants

23 native German speakers (22 female) with no self-reported hearing deficits were recruited from Maastricht University’s student population and gave their informed consent to participate in the experiment. Of these participants, 2 were excluded due to failure to perform the task adequately (mean accuracy < 60%), and 1 was excluded due to technical issues during the recording resulting in fewer than 8 usable experimental blocks (i.e., 4 minutes of speech tracking data per exposure and condition). This resulted in a final sample of 20 participants. This study was carried out in accordance with the Declaration of Helsinki and approved by the ethical committee of the Faculty of Psychology and Neuroscience at Maastricht University (ERCPN_167_09_05_2016).

### 2.2 Procedure

#### 2.2.1 Stimuli and Task

Stimuli consisted of the first 6 min of a German audiobook (‘Das Nebelhaus’ by Eric Berg) spoken by a female speaker (F0 = 159 +/- 8.3 Hz), split into twelve 30-s excerpts. In twelve experimental blocks, participants were presented with one excerpt of the original speech or of single-channel noise vocoded speech, referred to as *intelligible* and *unintelligible* speech, respectively. The same 30-s excerpt was presented 5 consecutive times in each condition (i.e., intelligible or unintelligible), with the order of conditions alternating across blocks (Figure 1). We refer to these each consecutive stimulus presentation as “exposure”, abbreviated E1 – E5. To monitor their attention across the experiment, participants were instructed to detect amplitude modulations (AMs) that could occur between 1-5 times in each 30-second trial, and report the number of detected AMs at the end of the trial by button press.

**Figure 1.**
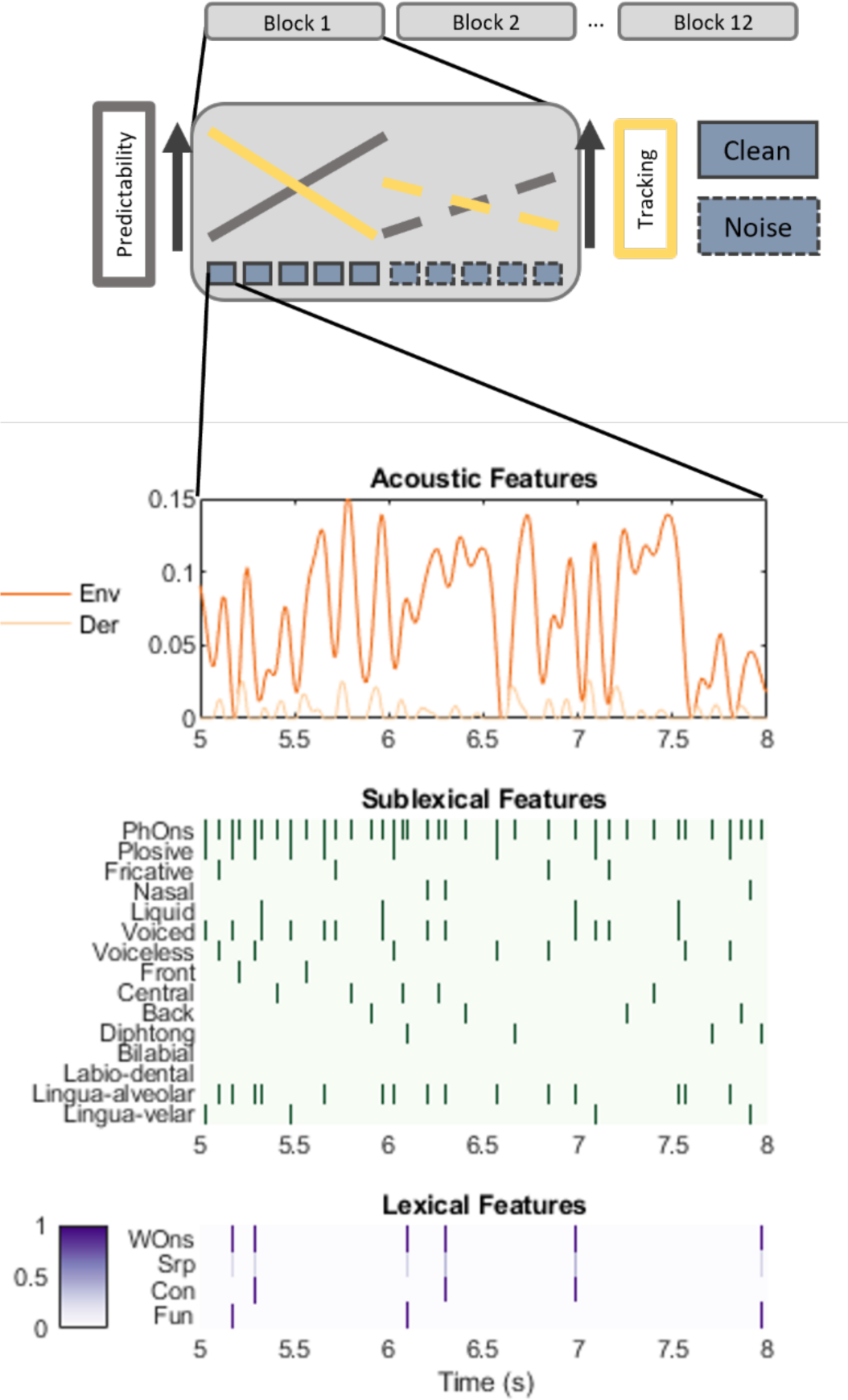
Overview of study design. The top panel illustrates the structure of each of the 12 experimental blocks and the main hypotheses. In each block, participants listened to five consecutive exposures of the same 30-s audiobook excerpt, presented either as intelligible or unintelligible speech, followed by five consecutive exposures in the other condition. The order of conditions alternated per block, and was counterbalanced across participants. We expected that the predictability of the input would increase with each exposure (grey bar), resulting in decreased speech tracking (yellow bar). We expected this pattern to be attenuated for unintelligible noise-vocoded speech (dashed lines) compared to clean intelligible speech (solid lines). The bottom panel illustrates the stimulus features included as predictors in the model: acoustic features consisted of the amplitude envelope (Env) and its derivative (Der); sublexical features consisted of phoneme onsets (PhOns) and 14 phonetic features, coded as binary predictors at phoneme onsets; Lexical features consisted of word onsets (WOns), GPT2-surprisal values (Srp), and parts of speech grouped as content (Con) and function (Fun) words, also coded as binary predictors at the word onsets.

#### 2.2.2 EEG data acquisition

64-channel EEG was acquired using the EasyCap montage 11 and BrainVision Recorder (Brain Products GmbH, Gilching, Germany), with a sampling rate of 500 Hz and band-pass filtered (cut-offs: 0.01 and 200 Hz, analog filter). The online reference was placed at position TP9 and the ground electrode at FT10. Two electrodes were placed below and beside the left eye to record vertical and horizontal eye movements, resulting in 62 scalp EEG channels and 2 EOG channels. The scalp was cleaned with skin preparation scrub and electrodes were filled with an electrolyte gel to keep impedances below 10 kOhm during the recording.

After preparing the EEG cap, participants inserted the in-ear earphones and were presented with two conditions to familiarize themselves with the stimuli and the task. The volume was initially set to 65 dB_SPL_ and then individually adjusted to a comfortable level. During the 6 practice trials (3 trials for each condition, i.e. clean and noise-vocoded speech), participants were given cues to when the AMs occurred during the first trial in each condition to ensure they could detect them accurately during the experiment.

During each experimental block, participants fixated on a fixation cross at the center of the screen, and were instructed to minimize blinking during the stimulus presentation. Each 30-s stimulus presentation was followed by a 1.5-s pause, during which participants were prompted to indicate how many AMs they counted, by pressing the appropriate number key (1 – 5). After each block, participants were encouraged to take a short break. Each participant completed 10 to 12 experimental blocks, depending on expired time and fatigue.

### 2.3 Analysis

#### 2.3.1 Behavior

Statistical analyses were performed in RStudio running R (version 4.5.0), using the libraries readxl, tidyr, ggpubr, ggplot2, rstatix. Due to the non-normal distribution of the accuracy data, non-parametric Friedman tests were used as an alternative to a repeated-measures ANOVA to test for an effect of Exposure (1 – 5) on accuracy scores (proportion of blocks where participants reported the correct number of amplitude modulations) separately for intelligible and unintelligible speech. To test for an interaction between Exposure (E1-E5) and Condition (intelligible and unintelligible speech), we calculated the difference in accuracy between the intelligible and unintelligible conditions for each exposure, and performed a Friedman test. Post-hoc contrasts were performed using paired Wilcoxon signed-rank tests, corrected for multiple comparisons using Bonferroni correction.

#### 2.3.2 EEG

##### 2.3.2.1 EEG preprocessing

EEG preprocessing was carried out in Matlab (v R2020a; The Mathworks, Natick, MA, United States) using the EEGLAB toolbox (v2021.0). The data were band-pass filtered between 0.5 Hz and 45 Hz to remove low-frequency drifts as well as high-frequency artifacts such as muscle activity and line noise, and subsequently downsampled to 100 Hz. A faulty channel FPz was identified in 17 participants and interpolated with spherical interpolation. Data were re-referenced to the average of all scalp electrodes, and filtered between 0.5 and 8.5 Hz.

##### 2.3.2.2 Encoding models

Increasingly complex forward (encoding) models were trained at three acoustic/linguistic processing levels, separately for intelligible and unintelligible speech: acoustic, sublexical, and lexical (Figure 1). The acoustic model included two acoustic predictors: amplitude envelope and its derivative. The sublexical model included both acoustic predictors, as well as a predictor for phoneme onset, and 14 phonetic features. Finally, the lexical model included all acoustic and sublexical predictors, as well as predictors for word onset, GPT2-surprisal, and parts of speech (content vs. function words).

###### Acoustic predictors

The acoustic predictors coded the speech signal’s amplitude envelope (Env) and its derivative (Der). The amplitude envelope was extracted from the speech signal using the absolute value of the Hilbert transform, downsampled to 100 Hz, and then low-pass filtered with a cut-off of 10 Hz. Values below 0.001 were set to 0. The derivative was calculated as the difference between two consecutive envelope values and rectified (i.e., with negative values set to 0). This predictor represents acoustic onsets in the speech signal.

###### Sublexical predictors

The sublexical predictors consisted of phoneme onsets (PhOns) and 14 phonetic features (PhonFeat). Phonemes were extracted from the continuous speech signal using the Munich Automatic Segmentation system (MAUS; Schiel, 1999) to automatically segment the speech signal into phonetic units and label these. These labels were subsequently manually reviewed and corrected by a native German speaker using Praat software (version 6.2.22, Boersma & Weenink, 2013), and then converted to Matlab matrices representing the onset of each phoneme at a 100 Hz sampling rate using the mPraat toolbox (Bořil & Skarnitzl, 2016). These phonemes were then converted to 19 phonetic features (Di Liberto et al., 2015; using categorization from the University of Iowa’s phonetics project https://soundsofspeech.uiowa.edu/german) describing manner of articulation (plosive, fricative, affricate, nasal, liquid, glide), place of articulation (bilabial, labio-dental, lingua-dental, lingua-alveolar, lingua-palatal, lingua-velar, glottal) and voicing (voiced, voiceless) for consonants, as well as backness (front, central, back) and diphthong for vowels. Due to fewer than 10 observations per block on average for glides, affricates, glottal, lingua-dental and lingua-palatal consonants, these five predictors were excluded from subsequent analyses. This resulted in a total of 15 sublexical predictors, including 1 predictor of phoneme onset (PhOns), marking the onset of a phoneme, and 14 phonetic features (PhonFeat): plosive, fricative, nasal, liquid (manner of articulation), bilabial, labio-dental, lingua-alveolar, lingua-velar (place of articulation), voiced, voiceless (voicing), front, central, back, diphthong (vowels).

###### Lexical predictors

Lexical predictors consisted of word onsets, GPT2-surprisal (e.g., Caucheteux et al., 2023), and parts of speech (POS; e.g., Gillis et al., 2021). Word onsets were parsed out from the output of the Munich Automatic Segmentation system (MAUS; Schiel, 1999), which reported the time of the word onsets based on the audio files of the audiobook excerpts. The reported times were then transformed into the timeseries of binary predictors.

The word surprisals were calculated using the German GPT-2 Large Language Model trained by the MDZ Digital Library team at Bavarian State Library (*Dbmdz/German-Gpt2 · Hugging Face*, n.d.). The model was hosted on the HuggingFace model hub. The surprisal scores were retrieved from the model using the minicones Python package (Misra, 2022). Minicones was designed to automate the token probability retrieval with respect to the trained language model parameters. Token in this context refers to the word unit that the large language model uses to learn the semantic relationship between tokens. Tokens are either the fragments of the word or the words themselves. The German GPT-2 model uses 50.000 unique tokens identified with the Byte-Pair Encoding algorithm during training. The surprisals were extracted in a sentence-by-sentence manner using an incremental language model scorer. The incremental language model scorer was used to extract the conditional probability of the token provided the preceding tokens in the sentence, as opposed to using both preceding and following tokens to identify the token probability. In a final step, the surprisal values for words were calculated as a sum of the log probabilities of the tokens comprising the word (joint probability; e.g., Heilbron et al., 2022).

To extract parts of speech information, the spaCy Python package for natural language processing was used (Montani et al., 2023). The spaCy package is designed to extract linguistic features using a pretrained pipeline. The “de_core_news_sm” pipeline was used, due to its high parts-of-speech labeling capability (self-reported 98% accuracy). The transcripts of the audiobook excerpts were labeled with Universal POS tags (i.e. noun, verb, pronoun). The dimensionality was reduced by categorizing parts of speech as Content and Function words, which resulted in two binary predictors.

##### 2.3.2.3 Forward modeling of EEG signals

EEG signals were modelled through regularized linear regression using multivariate temporal response functions as implemented in the mTRF toolbox (Crosse et al., 2016, 2021). To avoid processes related to acoustic onsets of stimuli influencing the modelling, the first 2 s of the trials were excluded from the analysis. Delays ranging from −0.1 s to 1.0 s relative to the predictors were considered, in order to capture both early acoustic and later linguistic processes.

In an initial cross-validation procedure using the mTRFcrossval.m function from the mTRF toolbox, an optimal lambda for regularization was selected from 14 possible values (λ = {10^−1.5^, 10^−1^,…, 10^0^,…, 10^4.5^, 10^5}^). For each participant, model type (acoustic, sublexical, lexical) and condition (clean vs. noise-vocoded speech), the best lambda value was determined, pooling across blocks and exposures. This was achieved through a leave-one out procedure, where the lambda with the best model performance trained on k-1 blocks was determined. This approach allowed optimizing model regularization per participant, model type and condition, and crucially, ensured comparability across exposures.

Models were trained using a leave-one-out cross-validation procedure, where training data consisted of data from k-1 blocks of a single exposure. The fitted model was then tested on data from each repetition of the left-out block. For example, a model trained on data of the first exposure of blocks 1 to 11 was tested on exposures 1 through 5 of block 12. Model performance was quantified as the linear correlation between the predicted and the actual EEG signal for the left-out data using Pearson’s correlation coefficient. As a measure of the noise ceiling for model performance, we calculated the consecutive exposure correlation. For each block, EEG activity from one exposure was correlated with the average of the other four exposures. This was repeated for each block and exposure, and the results were averaged per participant and condition.

##### 2.3.2.4 Statistical Analysis

Statistical analyses were performed in RStudio running R (version 4.5.0), using the libraries readxl, tidyr, ggpubr, ggplot2, rstatix. In a first step, we compared model performance between model types (acoustic, sublexical, lexical), averaged across blocks and exposures, to determine which model type best suited the data. Due to the presence of outliers, analyses were performed using non-parametric Friedman tests with model performance as the outcome variable, and model type (acoustic, sublexical, lexical) as a predictor. These analyses were performed separately for the clean speech and noise conditions. Post-hoc paired Wilcoxon signed-rank tests were Bonferroni corrected for multiple comparisons.

Subsequent analyses were performed on data from the sublexical model (i.e., models based on both acoustic and sublexical features). If increased predictability results in reduced neural encoding of the speech signal, we would expect model performance to decrease with each subsequent exposure. To assess whether the encoding of speech is affected by exposure, we compared model performance across exposures when models were trained and tested on the same exposure using non-parametric Friedman tests with model performance as the outcome variable, and exposure (E1 – E5) as the predictor. This analysis was performed separately for intelligible and unintelligible speech.

In additional exploratory analyses, we examined whether the neural encoding of the incoming speech signal was exposure-specific. We quantified ‘encoding specificity’ as the difference between model performances observed with left-out data from the exposure on which the model was trained vs. data from all other exposures. For example, for a model trained on exposure E1, we compared its performance on exposure E1 to its performance on exposures E2 to E5. A value of 0 would indicate that the model performance does not change when the model is tested on data from a different exposure than the exposure it was trained on. If the encoding is exposure-specific, we would expect exposure specificity to be significantly greater than 0, indicating higher model performance when the training and testing exposures are the same. This was tested using one-sided one-sample Wilcoxon signed rank tests for clean and noise-vocoded speech separately. We further performed a one-sided paired Wilcoxon signed rank test comparing encoding specificity between intelligible and unintelligible speech.

To verify whether an effect of repeated exposure on encoding specificity reflects predictive processes, we tested whether model performance changes with the temporal distance between the training and testing data. If an effect of repeated exposure is related to predictive processes, we expect increasing temporal distance to be associated with greater changes in model performance compared to the training model. To test this, we fit a linear mixed effects model with the difference in model performance as the outcome variable, and distance from the training data as a predictor, for intelligible and unintelligible speech separately.

Finally, we examine the time course of this putative distance effect in the TRFs encoding the envelope, its derivative, and phoneme onsets. The standardized TRF weights for speech envelope, envelope derivative, and the phoneme onsets, were separately submitted to a cluster-corrected permutation analysis (Maris & Oostenveld, 2007), comparing the first exposure to each subsequent exposure (E1 vs. E2, E1 vs. E3,…). In an additional analysis, consecutive exposures (E1 vs. E2, E2 vs. E3,…) were also compared.

## 3 Results

### 3.1 Behavior

Overall accuracy scores were high, with mean accuracy scores of over 80% across all conditions and exposures (Figure 2A). A non-parametric Friedman test examining the effect of exposure on the difference in accuracy between conditions revealed a significant result (X^2^ (4) = 16.39, p = 0.0025), suggesting an interaction between the variables condition and repetition (Figure 2B). Repeating the Friedman test using the accuracy scores for intelligible and unintelligible speech separately revealed a significant effect of exposure on accuracy for intelligible (X^2^(4) = 11.3, p = 0.0231), but not for unintelligible speech (X^2^(4) = 6.36, p = 0.174), suggesting that this interaction is driven by differences within the intelligible speech condition. Post-hoc paired Wilcoxon signed-rank tests comparing conditions for each exposure failed to reach significance after Bonferroni correction, however a trend for higher accuracy scores in the clean speech condition was observed for E1 (W = 152, p.adj = 0.0535) and E2 (W = 97, p.adj = 0.0935). Pairwise comparisons between exposures for intelligible speech also did not reveal any significant contrasts after Bonferroni correction, aside from a trend towards decreased accuracy for R4 compared to R1 (W = 130, p.adj = 0.095). We note that the highest accuracy was observed for exposure E1 (94.1%), compared to the other exposures with scores ranging between 84.7% (E3) and 89.1% (E2).

**Figure 2.**
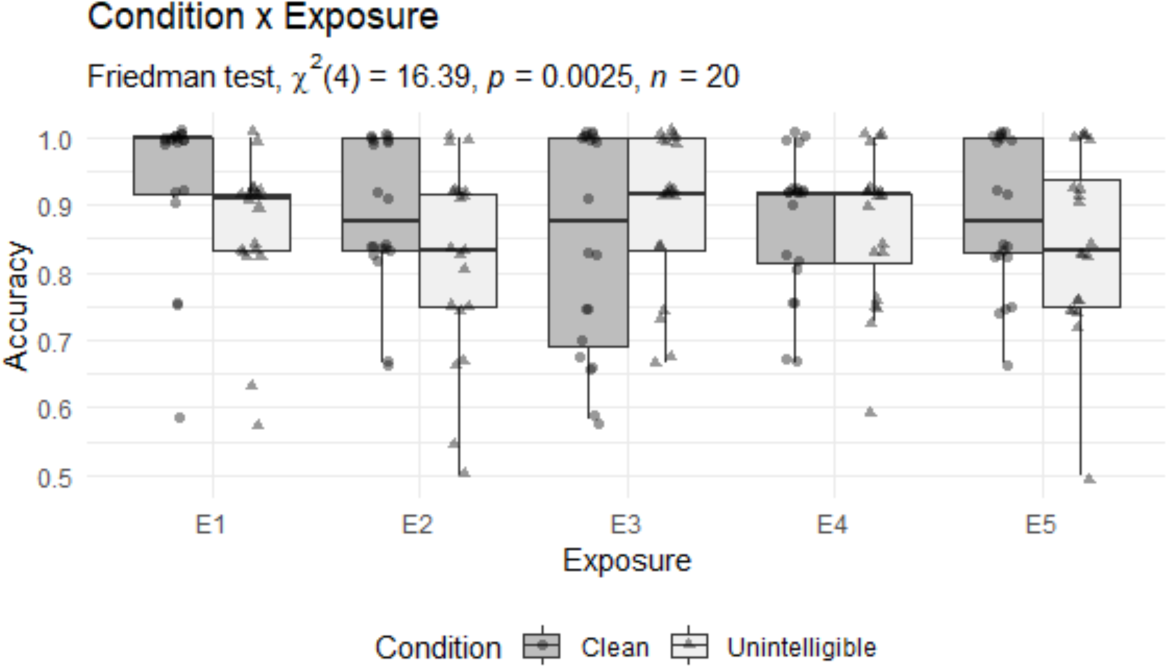
Behavioral results. Target detection accuracy plotted across exposures (E1 – E5) and conditions (clean vs. unintelligible speech). A non-parametric Friedman test on the difference between conditions revealed a significant interaction between Condition and Exposure on accuracy; however, no pairwise comparisons remained significant after Bonferroni correction.

### 3.2 Encoding models

#### 3.2.1 Model comparisons

Non-parametric Friedman tests were used to compare different model types (acoustic, sublexical and lexical; Figure 3A-C), averaged across blocks and exposures, separately for clean and noise-vocoded speech. These tests revealed differences in model performance across model types for both intelligible (X^2^= 6.3, p = 0.043) and unintelligible speech (X^2^= 12.7, p = 0.0017; Figure 3D). Post-hoc paired Wilcoxon signed-rank tests (Bonferroni corrected) revealed that the inclusion of sublexical predictors (phoneme onset and phonetic features) significantly improved model performance compared to an acoustic-only model for intelligible speech (W = 43, p.adj = 0.038), but that there was no added benefit of lexical predictors (GPT2-surprisal, parts of speech; W = 135, p.adj = 0.554). Interestingly, in the unintelligible condition including sublexical predictors worsened model performance (W = 188, p.adj = 0.002).

**Figure 3.**
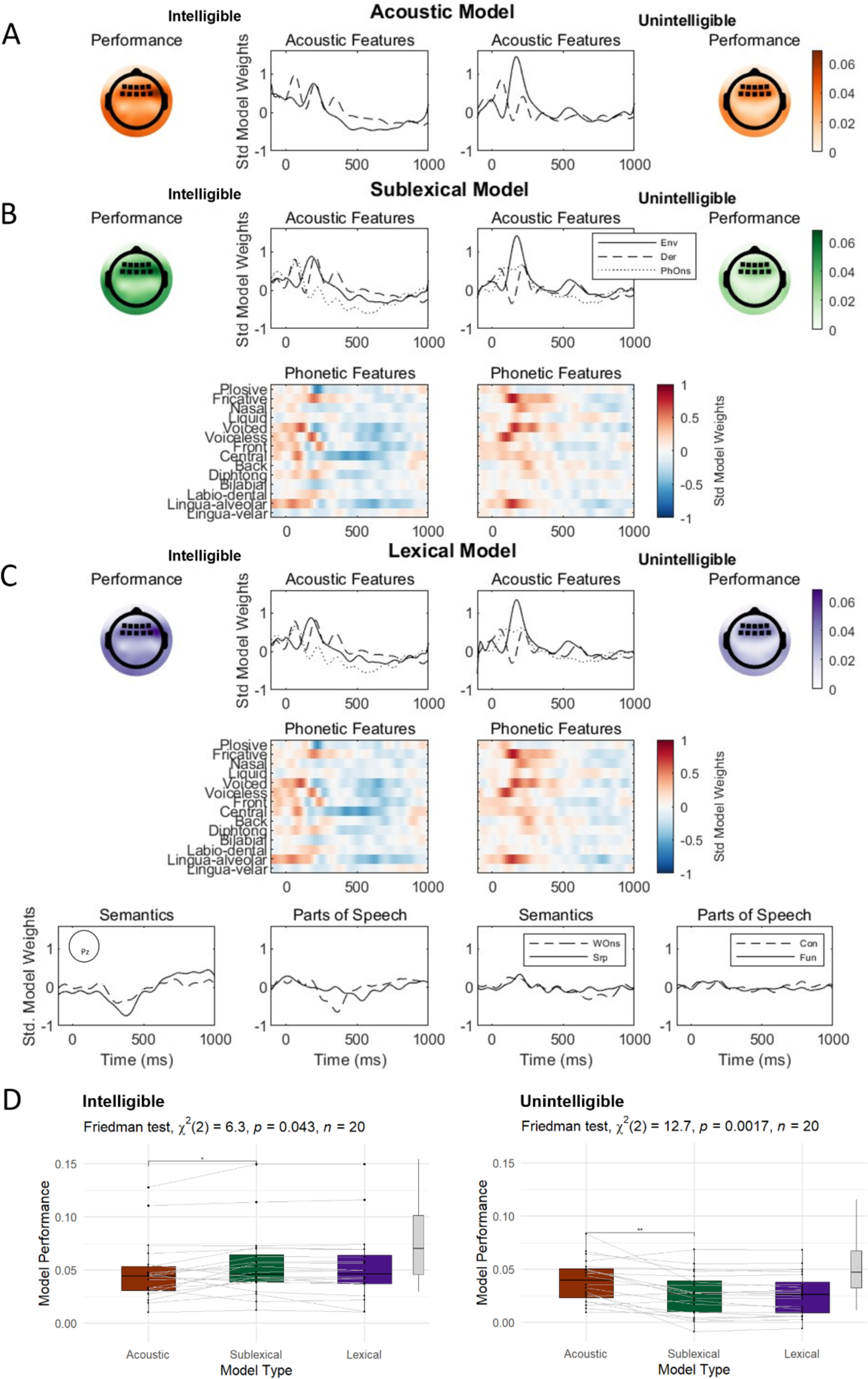
Overview of encoding models. Linear encoding models including predictors for different processing levels are presented for intelligible (left) and unintelligible speech (right). Topographies represent model performance averaged across all repetitions, TRFs for acoustic and sublexical predictors are averaged within a frontocentral ROI indicated in topographic maps. TRFs from lexical predictors are plotted at Pz to show N400-like pattern. (A) Acoustic model including predictors for envelope (Env, solid line) and envelope derivative (Env, dashed line). (B) Sublexical model including acoustic predictors as well as phoneme onsets (PhOns, dotted line) and 14 phonetic features. (C) Lexical model including all acoustic and sublexical features as well as word onsets (WOns, dashed line), GPT2 surprisal (Srp), and parts of speech coded as content (Con, dashed line) and function (Fun, solid line) words. (D) Model comparisons reveal that including sublexical features improves model fit over acoustic features for intelligible speech, but there is no added benefit of lexical features (left plot). For unintelligible speech, the acoustic model performs best (right plot). Grey boxplots on the right of each panel represents the consecutive exposure correlation as a measure of noise ceiling for model performance (see Methods 2.3.2.3).

These findings indicate that the sublexical model is the most suited for capturing neural processing of the speech signal in the current dataset. Thus, subsequent analyses are performed using this model.

#### 3.2.2 Repetition effect

Non-parametric Friedman tests were used to compare model performance across exposures when the sublexical models were trained and tested on the left-out block from the same exposure (Figure 4A, values on the diagonal). These analyses were performed separately for the intelligible and unintelligible condition, with model performance as the outcome variable, and exposure as the predictor variable. These tests revealed no effect of exposure on model performance for intelligible (X^2^(4) = 3.72, p = 0.45) or unintelligible speech (X^2^(4) = 2.6, p = 0.63; Figure 4B).

**Figure 4:**
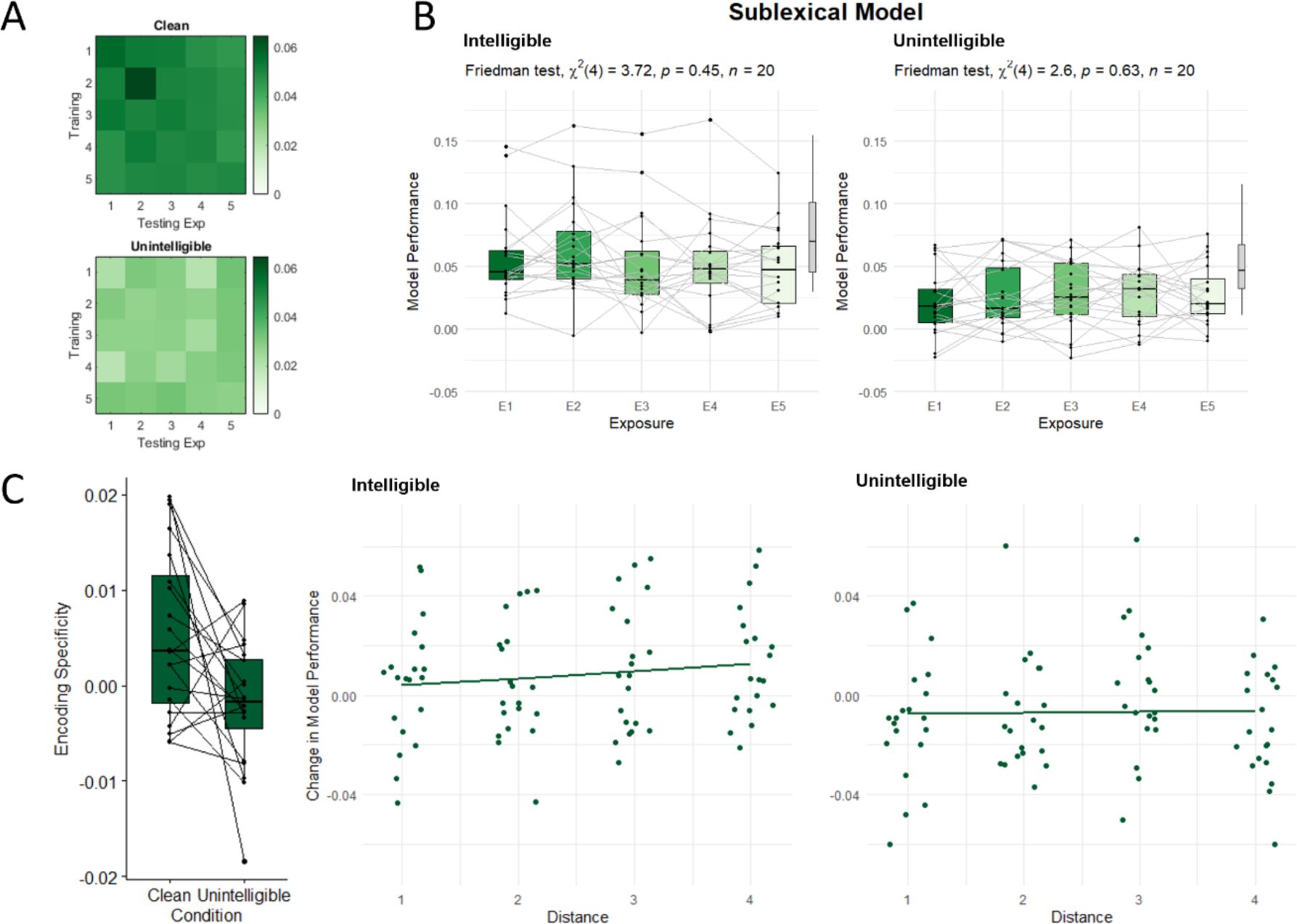
Model performance across exposures. (A) Average model performance, averaged across blocks and in a frontocentral ROI, for all combinations of training and testing exposures. (B) Individual model performance across exposures when the model was trained and tested on the same exposure (on-diagonal). Grey boxplots on the right of each panel represents the consecutive repetition correlation, a measure of the noise ceiling for model performance (see Methods 2.3.2.3). (C) Encoding specificity, calculated as the mean difference between on-diagonal (training exposure = testing exposure) and off-diagonal (training exposure ≠ testing exposure) model performance (left). Change in model performance with increasing temporal distance, for models trained on the first exposure and tested on all subsequent exposures, with distance being the difference between training and testing exposures (e.g., E2-E1 = 1, E3-E1 = 2, etc.).

#### 3.2.3 Encoding specificity

Encoding specificity, defined as the difference in model performance between the training exposure and the average of all other exposures, was tested in one-sided one-sample Wilcoxon signed rank tests for clean speech and noise separately. Encoding specificity was significantly greater than 0 for intelligible speech (W = 162, p = 0.0164), indicating that cortical speech tracking processes are exposure-specific, or conversely, do not generalize across exposures for intelligible speech. This was not the case for unintelligible speech (W = 78, p = 0.844), suggesting that this exposure specificity is contingent on the availability of linguistic information. A direct comparison between speech conditions confirmed that encoding specificity was significantly greater for intelligible compared to unintelligible speech (W = 170, p = 0.00681; Figure 4C).

With each repeated exposure to the same speech input, participants become more familiar with its content, and can form stronger predictions about the unfolding speech. If the observed encoding specificity is related to prediction processes, we would expect the difference in model performance to change with increasing distance from the training exposure. To test this, we investigated the change in model performance across exposures for models trained on the first exposure (Figure 4C). We calculated the difference in model performance between the first exposure and each subsequent exposure (e.g., E1 – E2, E1 – E3, etc.). These difference scores were used as the outcome variable in a linear mixed effects model, with the distance from the training exposure (e.g., for E1 – E2, the distance is 1, for E1 – E3, the distance is 2, etc.) as the outcome variable, and random intercepts per participant. For intelligible speech, this yielded a significant effect of distance (β= 0.002879, std. error = 0.001252, p = 0.0251), with increasing distance from the training exposure resulting in a larger decrease in model performance. The same analysis was repeated for the unintelligible speech condition, where no effect of distance was observed (β= 0.000455, std. error = 0.001343, p = 0.736), suggesting that this effect is specific to the availability of linguistic information, and not simply related to other factors related to the passage of time.

To further support that this encoding specificity effect is related to linguistic predictions, these analyses were repeated with the acoustic model. Similar to the sublexical models, the acoustic models showed significant encoding specificity for intelligible (W = 159, p = 0.022), but not for unintelligible speech (W = 116, p = 0.351). However, a direct comparison did not reveal a significant difference in model specificity between the two conditions (W = 140, p = 0.101), suggesting that this effect is less robust for the acoustic model. A linear mixed effects model did not reveal an effect of distance on the change in model performance for intelligible (β= 0.001677, std. error = 0.001622, p = 0.305) or unintelligible speech (β= 0.000782, std. error = 0.00124, p = 0.529).

One-sided *z*-tests were performed to compare the slopes of the distance effects. While the comparison between sublexical and acoustic models for intelligible speech did not yield a significant result (z = 0.5866, p = 0.279), we observed a statistical trend towards a difference in the slopes between intelligible and unintelligible speech for the sublexical model (z = 1.320, p = 0.0934). Taken together, this evidence suggests that linguistic (sublexical) predictions have a greater impact on the neural encoding of the speech signal compared to acoustic predictions, only in intelligible speech.

#### 3.2.4 Temporal response functions

For the intelligible speech condition, the standardized TRF weights from the sublexical model for speech envelope, envelope derivative, and the phoneme onsets, averaged within a frontocentral ROI, were separately submitted to a cluster-corrected permutation analysis, comparing the first exposure to each subsequent exposure (E1 vs. E2, E1 vs. E3,…;Figure 5). The additional 14 phonetic features were excluded from this analysis as phoneme-specific processing differences are beyond the scope of this analysis. For the speech envelope, this analysis identified a significant cluster for the contrast E1 vs. E5, for the time points −90 ms to −10 ms relative to the speech envelope (p = 0.035). Here, increased model weights for the fifth exposure (E5)_compared to the first exposure (E1) may suggest predictive encoding of the unfolding signal. For the envelope derivative, significant clusters were identified for the contrasts E1 vs. E3 at delays 40 ms –130 ms relative to the derivative (p = 0.012), and E1 vs. E4 at delays 70 ms –130 ms relative to the derivative (p = 0.032). These differences were all characterized by larger absolute model weights during later exposures, potentially indicating a predictive enhancement of the neural encoding of the speech signal. No significant clusters were identified for the phoneme onsets, or for the comparisons of consecutive exposures (e.g., E2 vs. E3, etc.). An overview of the topographic distribution of the model weights within these clusters can be observed in Supplementary Figure S1.

**Figure 5:**
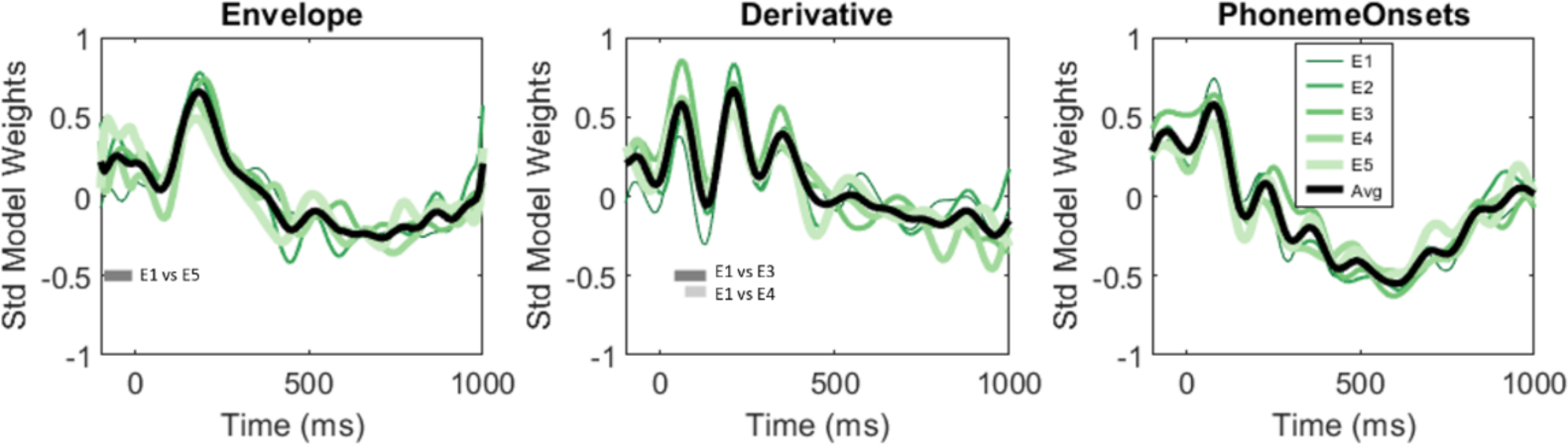
Changes in temporal response functions across exposures for intelligible speech. Standardized TRFs per exposure for speech envelope, its derivative, and phoneme onsets. Significant clusters highlighted via grey horizontal lines.

## 4 Discussion

The current experiment aimed to study the effect of predictability of the speech signal on cortical speech tracking. We operationalized predictability by repeatedly exposing participants to the same 30-s speech signal, presented either as intelligible clean speech or unintelligible single-channel noise-vocoded speech. Participants counted amplitude modulations occurring 1-5 times, and performed this task with high accuracy across both conditions, with a trend towards higher performance in the clean speech condition for the first two exposures. In our EEG analyses, we did not observe the hypothesized changes in model performance across exposures, when training and testing exposure were the same. This indicates that increased predictability, when operationalized as repeated exposure to the same stimulus, may not decrease (or increase) neural encoding of that signal. Instead, exploratory analyses revealed that cortical encoding was exposure-specific, meaning that models performed better when they were trained and tested on the same exposure, compared to when they are tested on a different exposure than the training exposure. The difference in model performance increased with increasing distance from the training exposure, an observation we made only for clean speech. This suggests that the effect is due to increased familiarity with the linguistic signal. Looking into the time course of these changes in neural encoding, analyses revealed differences in TRFs in early time windows, indicative of early predictive enhancement of speech processing. Taken together, these findings suggest that linguistic predictability modulates cortical tracking of sublexical speech features.

Prevailing theories of predictive processing propose reduced neural processing for more predictable sensory input (Aitchison & Lengyel, 2017; Friston, 2005; Rao & Ballard, 1999; Spratling, 2017). Such observations have been reported in conventional ERP analyses of predictive language processing across linguistic processing levels (e.g., Nour Eddine et al., 2024; Ylinen et al., 2016). Recently, similar patterns have been observed for neural encoding of continuous speech (Schubert, Schmidt, et al.) and music (Keitel et al., 2025) envelopes. In line with these observations, in the current design we would expect the neural encoding of the speech signal, as indexed by model performance, to decrease with repeated exposure. However, we did not observe any changes in model performance across exposures when the models were trained and tested on the same exposure (Figure 4B). This deviation from the previous studies may be explained by differences in how “predictability” is operationalized across experimental designs. Schubert, Schmidt and colleagues manipulate predictability by introducing semantic violations in their stimuli, similarly to a classical N400 paradigm. Keitel and colleagues differentiate between high and low pitch predictability in melodies, based on (Western) tonal principles associated with certain pitch organizations allowing higher predictability of sequences compared to atonal music (Mencke et al., 2019; Ockelford & Sergeant, 2013). In contrast, the current design manipulates predictability by exposing participants to the same 30-s audiobook excerpt in five consecutive repetitions. Thus, the input signal is natural speech that does not contain clear violations or errors. The inherent predictability of the stimulus does not change, while the participants’ familiarity increases.

While our analyses did not show any change in *how well* the brain encodes the speech signal after repeated exposure, our exploratory analyses showed that neural encoding of the speech signal is nevertheless exposure-specific, meaning that model performance is highest when the models were trained and tested on the same exposure (Figure 4C). This suggests that the amount of prior knowledge about the incoming signal shapes how this signal is encoded. The difference in model performance was further shown to increase with increasing temporal distance between the training and testing exposure, in line with the idea that increasing familiarity with the input can lead to increased predictions. Crucially, no such effect of temporal distance was observed for noise-vocoded speech, or for the acoustic models, supporting that this effect relies on the availability—and operates on the encoding—of linguistic information, and is not merely due to external factors related to the passage of time (e.g., change in EEG signal quality participant movement, or fatigue).

To examine the time course of these changes in neural encoding, we analyzed changes in the TRFs across exposures. Here, we observed modulations in early time-windows for the TRFs representing the speech envelope and its derivative characterized by larger absolute model weights during later exposures in time lags preceding the stimulus envelope as well as aligned with typical P1/N1 latencies. This increased weighting of the signal might indicate a predictive enhancement of neural processing. This pattern stands in contrast to a recent study that showed reduced TRF amplitudes after repeated exposure at the P2 processing stage (Sánchez-Costa, Carboni, & Cervantes Constantino, 2025). These deviating results may be explained by different underlying predictive processes as proposed by Press and colleagues (2020), who propose a two-process model wherein perception is initially biased to expected input, resulting in early enhanced processing of predictable stimuli. Later processes may then reflect the prediction error generated by unexpected input, necessary for updating the internal model to improve future predictions. Sánchez-Costa and colleagues (2025) compared trials where the encoded speech signal was preceded by either the same speech segment or a different utterance. As there were twice as many trials where the speech signal was repeated compared to trials without repetition, this may have led to a “violation” response in trials without repetition, thus triggering enhanced encoding at the later P2 stage. Without any such violations in the current experiment, perceptual enhancement of the increasingly predictable input in early (−90 ms to 130 ms) time windows is in line with Bayesian theories and the two-process model of predictive processing.

An interesting observation in the current dataset is the fact that lexical predictors did not significantly improve model fit compared to the model containing acoustic and sublexical predictors. This stands in contrast to previous studies, which observed improved model performance for models including lexical surprisal (e.g., Broderick et al., 2018; Gillis et al., 2021; Heilbron et al., 2022). In the current design, participants were instructed to count amplitude modulations in the signal. While the choice of task ensured comparability across conditions (clean vs. noise-vocoded speech), it did not require participants to attend to the semantic level of the speech signal, which likely influenced how well the semantic information was encoded. Attention is known to influence cortical speech tracking (Carta et al., 2025; Hausfeld et al., 2018; O’Sullivan et al., 2015), and attention to specific features of the speech signal may modulate how those features are specifically tracked. Several studies have shown reduced tracking of linguistic features such as semantic surprisal (Dou et al., 2025) or phonetic features (Teoh et al., 2022) in unattended speech. In our design, the sublexical predictors may be more closely linked to the acoustic features the participants were attending to when detecting amplitude modulations, which may explain why sublexical features are still tracked despite not being explicitly attended to.

Taken together, our results indicate that repeated exposure to the same continuous speech input does not affect *how well* listeners track the speech input; however, increased familiarity with an unfolding speech signal shapes *how* listeners encode it. These differences arise in early time-windows (−90 to 130 ms), indicating early predictive enhancement of the unfolding signal. Crucially, these patterns are contingent on the availability of linguistic information: neither noise-vocoded, unintelligible speech nor models including only acoustic predictors showed the same patterns of encoding-specificity. As our current results are based on models including only acoustic and sublexical predictors, future research may consider investigating how these observations hold up for models including lexical predictors when listeners are attending to the semantic level of the speech input.

## Acknowledgements

The authors would like to thank Rosie Coppieters and Ahmad Jibai for their support in data collection. This research was supported by Maastricht University and the Netherlands Organisation for Scientific Research (NWO; VENI grant 451-17-033 to L.H.).

## Data availability

The data that support the findings of this study will be made openly available in Dataverse.nl upon acceptance.

## Author contributions

**Alexandra K. Emmendorfer:** Data curation, Formal analysis, Visualization, Writing – First version, Review & editing, Software, Methodology; **Lars Riecke:** Conceptualization, Writing – review & editing; **Hendrik Kröger:** Data curation, Resources, Writing – First version, Review & Editing; **Elizaveta Zavialova:** Data curation, Resources, Software, Writing – First version, Review & Editing; **Lars Hausfeld:** Conceptualization, Supervision, Writing – Review & editing, Funding acquisition, Methodology, Project administration, Investigation, Software, Resources

## Supplementary Materials

**Supplementary Figure S1:**
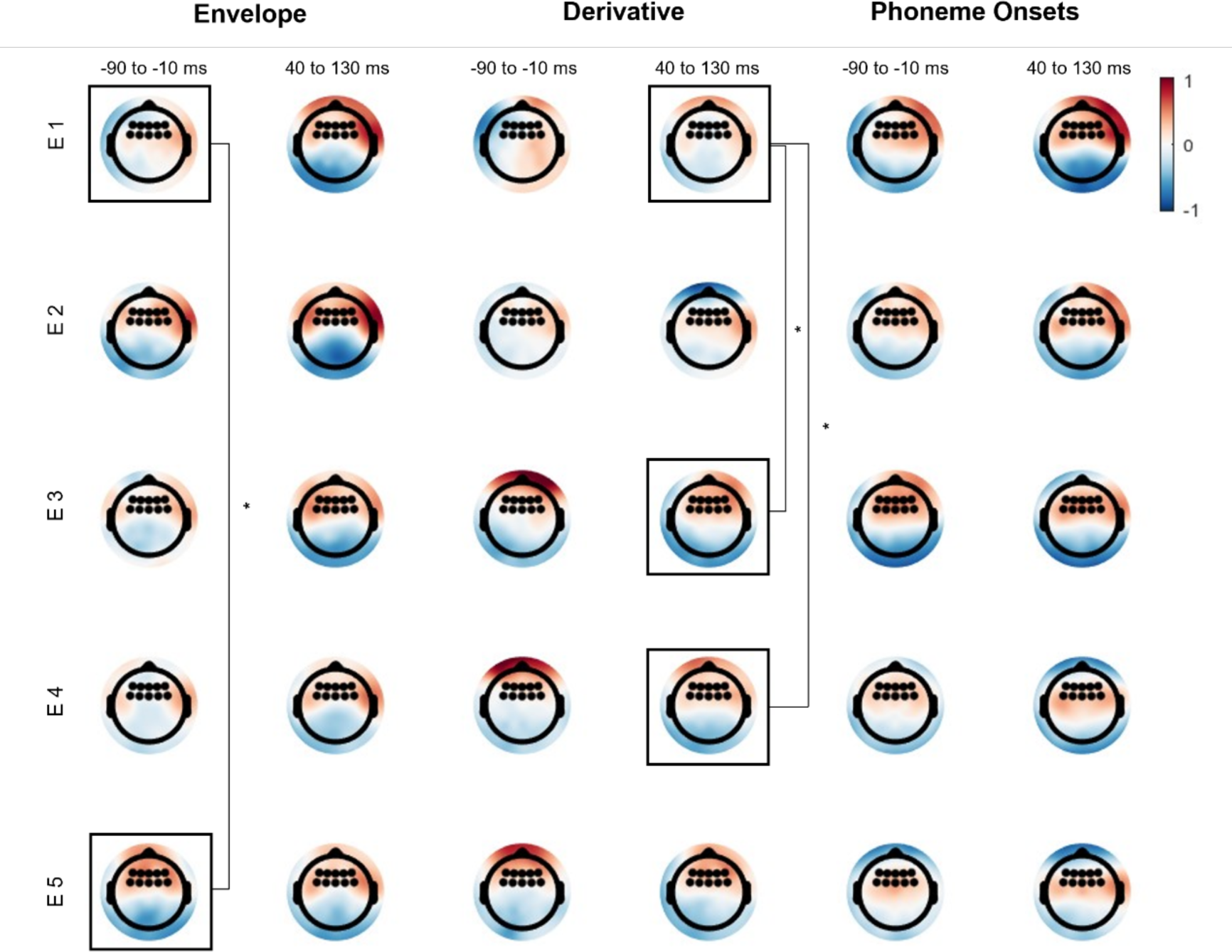
Topographic distribution of standardized model weights for sublexical predictors. Model weights are plotted averaged within time windows that were identified as significant clusters for the amplitude envelope (−90 to −10 ms), and its derivative (40 to 130 ms). Note: the contrast E1 vs E3 revealed a significant cluster for 40 to 130 ms, while the contrast E1 vs E4 revealed a significant cluster for 70 to 130 ms. For simplicity, only the window 40 to 130 ms is plotted.

## References

Aitchison, L., & Lengyel, M. (2017). With or without you: Predictive coding and Bayesian inference in the brain. Current Opinion in Neurobiology, 46, 219–227. 10.1016/j.conb.2017.08.010

Barthel, M., Tomasello, R., & Liu, M. (2024). Conditionals in context: Brain signatures of prediction in discourse processing. Cognition, 242, 105635. 10.1016/j.cognition.2023.105635

Boersma, P., & Weenink, D. (2013). Praat: Doing phonetics by computer. Version 5.3. 55. Http.

Bohn, K., Knaus, J., Wiese, R., & Domahs, U. (2013). The influence of rhythmic (ir)regularities on speech processing: Evidence from an ERP study on German phrases. Neuropsychologia, 51(4), 760–771. 10.1016/j.neuropsychologia.2013.01.006

Bonte, M. L., Mitterer, H., Zellagui, N., Poelmans, H., & Blomert, L. (2005). Auditory cortical tuning to statistical regularities in phonology. Clinical Neurophysiology, 116(12), 2765–2774. 10.1016/j.clinph.2005.08.012

Bořil, T., & Skarnitzl, R. (2016). Tools rPraat and mPraat. In P. Sojka, A. Horák, I. Kopeček, & K. Pala (Eds.), Text, Speech, and Dialogue (pp. 367–374). Springer International Publishing. 10.1007/978-3-319-45510-5_42

Brodbeck, C., Hong, L. E., & Simon, J. Z. (2018). Rapid Transformation from Auditory to Linguistic Representations of Continuous Speech. Current Biology, 28(24), 3976–3983.e5. 10.1016/j.cub.2018.10.042

Brodbeck, C., Kandylaki, K. D., & Scharenborg, O. (2024). Neural Representations of Non-native Speech Reflect Proficiency and Interference from Native Language Knowledge. Journal of Neuroscience, 44(1). 10.1523/JNEUROSCI.0666-23.2023

Brodbeck, C., & Simon, J. Z. (2020). Continuous speech processing. Current Opinion in Physiology, 18, 25–31. 10.1016/j.cophys.2020.07.014

Broderick, M. P., Anderson, A. J., Di Liberto, G. M., Crosse, M. J., & Lalor, E. C. (2018). Electrophysiological Correlates of Semantic Dissimilarity Reflect the Comprehension of Natural, Narrative Speech. Current Biology, 28(5), 803–809.e3. 10.1016/j.cub.2018.01.080

Carta, S., Aličković, E., Zaar, J., Valdés, A. L., & Di Liberto, G. M. (2025). Simultaneous cortical tracking of competing speech streams during attention switching. Neuroscience. 10.1101/2025.07.02.662762

Caucheteux, C., Gramfort, A., & King, J.-R. (2023). Evidence of a predictive coding hierarchy in the human brain listening to speech. Nature Human Behaviour, 7(3), 430–441. 10.1038/s41562-022-01516-2

Chen, Y.-P., Schmidt, F., Keitel, A., Rösch, S., Hauswald, A., & Weisz, N. (2023). Speech intelligibility changes the temporal evolution of neural speech tracking. NeuroImage, 268, 119894. 10.1016/j.neuroimage.2023.119894

Crosse, M. J., Di Liberto, G. M., Bednar, A., & Lalor, E. C. (2016). The Multivariate Temporal Response Function (mTRF) Toolbox: A MATLAB Toolbox for Relating Neural Signals to Continuous Stimuli. Frontiers in Human Neuroscience, 10. 10.3389/fnhum.2016.00604

Crosse, M. J., Zuk, N. J., Di Liberto, G. M., Nidiffer, A. R., Molholm, S., & Lalor, E. C. (2021). Linear Modeling of Neurophysiological Responses to Speech and Other Continuous Stimuli: Methodological Considerations for Applied Research. Frontiers in Neuroscience, 15. 10.3389/fnins.2021.705621

Dbmdz/german-gpt2 · Hugging Face. (n.d.). Retrieved August 13, 2026, from https://huggingface.co/dbmdz/german-gpt2

Di Liberto, G. M., Crosse, M. J., & Lalor, E. C. (2018). Cortical Measures of Phoneme-Level Speech Encoding Correlate with the Perceived Clarity of Natural Speech. Eneuro, 5(2), ENEURO.0084-18.2018. 10.1523/ENEURO.0084-18.2018

Di Liberto, G. M., O’Sullivan, J. A., & Lalor, E. C. (2015). Low-Frequency Cortical Entrainment to Speech Reflects Phoneme-Level Processing. Current Biology, 25(19), 2457–2465. 10.1016/j.cub.2015.08.030

Dou, J., Anderson, A. J., White, A. S., Norman-Haignere, S. V., & Lalor, E. C. (2025). Dynamic modeling of EEG responses to natural speech reveals earlier processing of predictable words. PLOS Computational Biology, 21(4), e1013006. 10.1371/journal.pcbi.1013006

Emmendorfer, A. K., Bonte, M., Jansma, B. M., & Kotz, S. A. (2023). Sensitivity to syllable stress regularities in externally but not self-triggered speech in Dutch. European Journal of Neuroscience, 58(1), 2297–2314. 10.1111/ejn.16003

Emmendorfer, A. K., Correia, J. M., Jansma, B. M., Kotz, S. A., & Bonte, M. (2020). ERP mismatch response to phonological and temporal regularities in speech. Scientific Reports, 10(1), 9917. 10.1038/s41598-020-66824-x

Emmendorfer, A. K., Jansma, B. M., Kotz, S. A., & Bonte, M. (2025). Neurophysiological responses to phonological and temporal regularities in speech in dyslexic and typical readers. Clinical Neurophysiology, 176, 2110772. 10.1016/j.clinph.2025.2110772

Etard, O., & Reichenbach, T. (2019). Neural Speech Tracking in the Theta and in the Delta Frequency Band Differentially Encode Clarity and Comprehension of Speech in Noise. Journal of Neuroscience, 39(29), 5750–5759. 10.1523/JNEUROSCI.1828-18.2019

Ferreira, F., & Chantavarin, S. (2018). Integration and Prediction in Language Processing: A Synthesis of Old and New. Current Directions in Psychological Science, 27(6), 443–448. 10.1177/0963721418794491

Friston, K. (2005). A theory of cortical responses. Philosophical Transactions of the Royal Society of London. Series B, Biological Sciences, 360(1456), 815–836. 10.1098/rstb.2005.1622

Gillis, M., Vanthornhout, J., Simon, J. Z., Francart, T., & Brodbeck, C. (2021). Neural Markers of Speech Comprehension: Measuring EEG Tracking of Linguistic Speech Representations, Controlling the Speech Acoustics. The Journal of Neuroscience, 41(50), 10316–10329. 10.1523/JNEUROSCI.0812-21.2021

Gwilliams, L., King, J.-R., Marantz, A., & Poeppel, D. (2022). Neural dynamics of phoneme sequences reveal position-invariant code for content and order. Nature Communications, 13(1), 6606. 10.1038/s41467-022-34326-1

Hausfeld, L., Hamers, I.M.H. & Formisano, E. (2024) FMRI speech tracking in primary and non-primary auditory cortex while listening to noisy scenes. Communications Biology, 7, 1217. 10.1038/s42003-024-06913-z

Hausfeld, L., Riecke, L., Valente, G., & Formisano, E. (2018). Cortical tracking of multiple streams outside the focus of attention in naturalistic auditory scenes. NeuroImage, 181, 617–626. 10.1016/j.neuroimage.2018.07.052

Heilbron, M., Armeni, K., Schoffelen, J.-M., Hagoort, P., & de Lange, F. P. (2022). A hierarchy of linguistic predictions during natural language comprehension. Proceedings of the National Academy of Sciences, 119(32), e2201968119. 10.1073/pnas.2201968119

Issa, M. F., Khan, I., Ruzzoli, M., Molinaro, N., & Lizarazu, M. (2024). On the speech envelope in the cortical tracking of speech. NeuroImage, 297, 120675. 10.1016/j.neuroimage.2024.120675

Keitel, A., Pelofi, C., Guan, X., Watson, E., Wight, L., Allen, S., Mencke, I., Keitel, C., & Rimmele, J. (2025). Cortical and behavioral tracking of rhythm in music: Effects of pitch predictability, enjoyment, and expertise. Annals of the New York Academy of Sciences, 1546(1), 120–135. 10.1111/nyas.15315

Koskinen, M., Kurimo, M., Gross, J., Hyvärinen, A., & Hari, R. (2020). Brain activity reflects the predictability of word sequences in listened continuous speech. NeuroImage, 219, 116936. 10.1016/j.neuroimage.2020.116936

Kotz, S. A., Ravignani, A., & Fitch, W. T. (2018). The Evolution of Rhythm Processing. Trends in Cognitive Sciences, 22(10), 896–910. 10.1016/j.tics.2018.08.002

Kubanek, J., Brunner, P., Gunduz, A., Poeppel, D., & Schalk, G. (2013). The Tracking of Speech Envelope in the Human Cortex. PLoS ONE, 8(1), e53398. 10.1371/journal.pone.0053398

Lesenfants, D., Vanthornhout, J., Verschueren, E., Decruy, L., & Francart, T. (2019). Predicting individual speech intelligibility from the cortical tracking of acoustic- and phonetic-level speech representations. Hearing Research, 380, 1–9. 10.1016/j.heares.2019.05.006

Liberto, G. M. D., Nie, J., Yeaton, J., Khalighinejad, B., Shamma, S. A., & Mesgarani, N. (2021). Neural representation of linguistic feature hierarchy reflects second-language proficiency. NeuroImage, 227, 117586. 10.1016/j.neuroimage.2020.117586

Maris, E., & Oostenveld, R. (2007). Nonparametric statistical testing of EEG- and MEG-data. Journal of Neuroscience Methods, 164(1), 177–190. 10.1016/j.jneumeth.2007.03.024

Mencke, I., Omigie, D., Wald-Fuhrmann, M., & Brattico, E. (2019). Atonal Music: Can Uncertainty Lead to Pleasure? Frontiers in Neuroscience, 12, 979. 10.3389/fnins.2018.00979

Misra, K. (2022). *minicons: Enabling Flexible Behavioral and Representational Analyses of Transformer Language Models* (arXiv:2203.13112). arXiv. 10.48550/arXiv.2203.13112

Montani, I., Honnibal, M., Honnibal, M., Boyd, A., Landeghem, S. V., & Peters, H. (2023). explosion/spaCy: V3.7.2: Fixes for APIs and requirements. 10.5281/zenodo.10009823

Noordenbos, M. W., Segers, E., Mitterer, H., Serniclaes, W., & Verhoeven, L. (2013). Deviant neural processing of phonotactic probabilities in adults with dyslexia. Neuroreport, 24(13), 746–750.

Norris, D., McQueen, J. M., & Cutler, A. (2016). Prediction, Bayesian inference and feedback in speech recognition. Language, Cognition and Neuroscience, 31(1), 4–18. 10.1080/23273798.2015.1081703

Nour Eddine, S., Brothers, T., Wang, L., Spratling, M., & Kuperberg, G. R. (2024). A predictive coding model of the N400. Cognition, 246, 105755. 10.1016/j.cognition.2024.105755

Ockelford, A., & Sergeant, D. (2013). Musical expectancy in atonal contexts: Musicians’ perception of “antistructure.” Psychology of Music, 41(2), 139–174. 10.1177/0305735612442582

O’Sullivan, J. A., Power, A. J., Mesgarani, N., Rajaram, S., Foxe, J. J., Shinn-Cunningham, B. G., Slaney, M., Shamma, S. A., & Lalor, E. C. (2015). Attentional Selection in a Cocktail Party Environment Can Be Decoded from Single-Trial EEG. Cerebral Cortex, 25(7), 1697–1706. 10.1093/cercor/bht355

Pickering, M. J., & Gambi, C. (2018). Predicting while comprehending language: A theory and review. Psychological Bulletin, 144(10), 1002–1044. 10.1037/bul0000158

Rao, R. P., & Ballard, D. H. (1999). Predictive coding in the visual cortex: A functional interpretation of some extra-classical receptive-field effects. Nature Neuroscience, 2(1), 79–87. 10.1038/4580

Schiel, F. (1999). Automatic Phonetic Transcription of Non-Prompted Speech.

Schubert, J., Schmidt, F., Gehmacher, Q., Bresgen, A., & Weisz, N. (2023). Cortical speech tracking is related to individual prediction tendencies. Cerebral Cortex, 33(11), 6608–6619. 10.1093/cercor/bhac528

Spratling, M. W. (2017). A review of predictive coding algorithms. Brain and Cognition, 112, 92–97. 10.1016/j.bandc.2015.11.003

Teoh, E. S., Ahmed, F., & Lalor, E. C. (2022). Attention Differentially Affects Acoustic and Phonetic Feature Encoding in a Multispeaker Environment. The Journal of Neuroscience, 42(4), 682–691. 10.1523/JNEUROSCI.1455-20.2021

Ter Bekke, M., Drijvers, L., & Holler, J. (2025). Co-Speech Hand Gestures Are Used to Predict Upcoming Meaning. Psychological Science, 36(4), 237–248. 10.1177/09567976251331041

Trujillo, J., Straube, B., & He, Y. (2025). Discourse context and co-speech gestures jointly shape hierarchical prediction during the processing of a multimodal narrative. PsyArXiv. 10.31234/osf.io/89qcn_v1

Vanthornhout, J., Decruy, L., Wouters, J., Simon, J. Z., & Francart, T. (2018). Speech Intelligibility Predicted from Neural Entrainment of the Speech Envelope. Journal of the Association for Research in Otolaryngology, 19(2), 181–191. 10.1007/s10162-018-0654-z

Vidal, Y., Brusini, P., Bonfieni, M., Mehler, J., & Bekinschtein, T. A. (2019). Neural Signal to Violations of Abstract Rules Using Speech-Like Stimuli. Eneuro, 6(5), ENEURO.0128-19.2019. 10.1523/ENEURO.0128-19.2019

Weissbart, H., Kandylaki, K. D., & Reichenbach, T. (2020). Cortical Tracking of Surprisal during Continuous Speech Comprehension. Journal of Cognitive Neuroscience, 32(1), 155–166. 10.1162/jocn_a_01467

Ylinen, S., Huuskonen, M., Mikkola, K., Saure, E., Sinkkonen, T., & Paavilainen, P. (2016). Predictive coding of phonological rules in auditory cortex: A mismatch negativity study. Brain and Language, 162, 72–80. 10.1016/j.bandl.2016.08.007

